# LDSC regression-based heritability estimates can be biased when summary statistics are obtained from meta-analysis or imputed variants

**DOI:** 10.64898/2026.07.05.736573

**Authors:** Rui Dong, Minghui Wang, Gao T. Wang, Andrew T. DeWan, Suzanne M. Leal

## Abstract

**Motivation:** Linkage disequilibrium score (LDSC) regression is a popular method to estimate heritability for complex traits using summary statistics and linkage disequilibrium (LD) reference panels, offering a practical alternative to methods requiring individual-level data. Despite its widespread use, LDSC regression can produce biased heritability estimates. The properties of LDSC regression were investigated using summary statistics from several large-scale Alzheimer’s disease (AD) studies and a variety of LD reference panels. These heritability estimates were compared with those obtained from individual-level data.

**Results:** When LDSC regression was applied to summary statistics obtained from meta-analysis, it led to an underestimation of heritability. This can occur if meta-analysis is used to combine studies of different ancestries leading to the caveat of the lack of an appropriate LD reference panel. Additionally meta-analyses often include studies with different phenotype definitions, that not only impacts heritability estimates but also makes them uninterpretable. Summary statistics generated from imputed variants, even those with high imputation accuracy, can lead to underestimation of heritability. For example, the heritability estimates for AD were reduced from 0.265 (se 0.148) to 0.160 (se 0.041) when imputed variants (INFO>0.9) were included compared to analyzing only genotype array variants. A decrease in heritability estimates was also observed when individual-level imputed variant data were analyzed using GCTA-GREML. Our findings highlight the caveats of estimating heritability using meta-analysis summary statistics or imputed data instead of genotyped or sequence data.

## Introduction

For the estimation of heritability using summary statistics obtained from genome-wide association studies (GWAS), Linkage Disequilibrium Score (LDSC) regression (Bulik-Sullivan *et al*. 2015) has been widely used. Unlike methods such as Genome-wide Complex Trait Analysis (GCTA)-Genomic-relatedness-matrix REstricted Maximum Likelihood (GREML) (Yang *et al*. 2011) that require individual-level data, LDSC regression provides a practical alternative, especially when access to individual-level data are restricted. Additionally for biobank-scale data it can be computationally prohibitive to estimate heritability using individual-level data. Despite the development of other methods to estimate heritability using summary statistics, e.g., Popcorn (Brown *et al*. 2016), GWASH (Schwartzman *et al*. 2019), HESS (Shi, Kichaev, and Pasaniuc 2016), and SumHer (Speed and Balding 2019), LDSC regression remains the preferred choice due to its straightforward implementation. LDSC regression has been used to estimate the heritability for a wide-variety of complex traits including Alzheimer’s disease (AD) (Wightman *et al*. 2021), type 2 diabetes (Jang *et al*. 2021), peripheral artery disease (Zuydam van *et al*. 2021), peptic ulcer disease (Wu *et al*. 2021), autism spectrum disorder (Grove *et al*. 2019), bipolar disorder (Mullins *et al*. 2021), etc (Sakaue *et al*. 2021).

There are discrepancies between narrow-sense heritability (h^2^) estimates derived from genetic data and those obtained from twin studies which are typically broad-sense heritability (H^2^) estimates. Unlike H^2^, h^2^ only includes additive genetic effects, therefore factors such as interaction will not contribute to the estimates. Additionally, usually only common single nucleotide variants (SNV) are used to estimate h^2^, although rare and structural variants may also play a role. For AD, twin studies consistently estimate H^2^ to be 0.6-0.8 (Bergem, Engedal, and Kringlen 1997, Gatz *et al*. 1997), while AD h^2^ estimates obtained from individual-level data range from 0.38-0.66 (Escott-Price and Hardy 2022, Baker *et al*. 2023). In contrast, recent large-scale GWAS (Alzheimer Disease Genetics Consortium (ADGC), *et al*. 2019, Jansen *et al*. 2019, Wightman *et al*. 2021, Bellenguez *et al*. 2022) that estimate h^2^ using LDSC regression, report much lower h^2^ estimates for AD, ranging from 0.018-0.106 (Table 1). These findings suggest substantial inconsistencies in h^2^ estimates depending on the method used and/or study design.

**Table 1.**
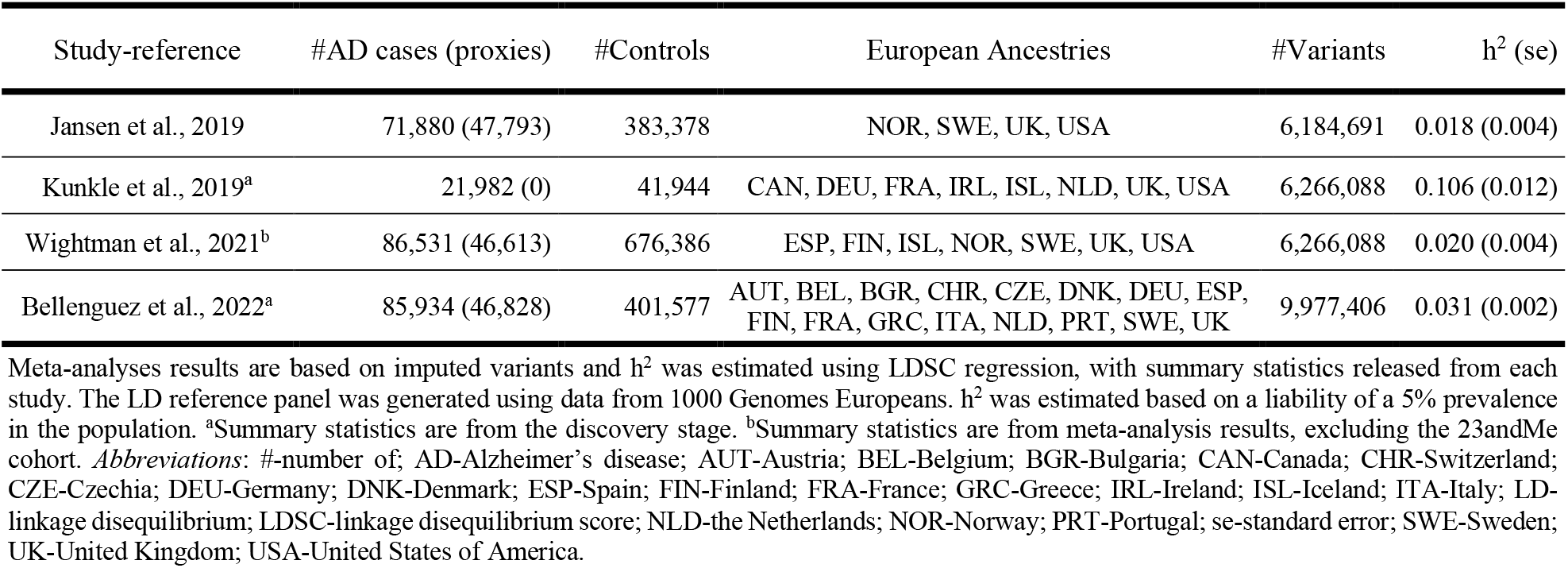
Alzheimer’s disease h^2^ estimates using meta-analysis results from large-scale studies.

These discrepancies are not isolated to AD; similar issues are observed for various complex traits (Jang *et al*. 2021, Mullins *et al*. 2021), raising broader concerns about the reliability of h^2^ estimates obtained from summary statistics. In this study, we examine several factors contributing to the limitations of estimating h^2^ using summary statistics through application of LDSC regression. Issues addressed are discrepancies between the ancestry of the study subjects used to obtain summary statistics and LD reference panels which can be more pronounced for meta-analyses, variations in phenotype definitions across studies, and the impact of analyzing summary statistics obtained from imputed variants. Using AD as a case study, we demonstrate how these factors can result in inconsistent and potentially misleading h^2^ estimates. Our analysis underscores the need for careful consideration of these factors to improve the accuracy and reliability of h^2^ estimates derived from summary statistics using LDSC regression.

## Materials and methods

White British UK Biobank AD cases and controls were identified using ICD10/ICD9 codes and self-report. AD cases (N=3,720) were defined as individuals with a primary or secondary diagnosis or cause of death including “AD” or “dementia in AD”. White British UK Biobank AD proxy cases (N=51,080) were defined as individuals without AD but have at least one parent or sibling with AD. There are 14,378 controls who are individuals >70 years-of-age without any record of AD or dementia-related illnesses and do not report having a family history of AD, i.e., parents or siblings. Since it was previously reported for LDSC regression that the analysis of summary statistics that include related individuals can create a bias (Lee *et al*. 2018), the analysis was limited to unrelated individuals. Related individuals (kinship coefficient >0.044) were identified and only one member of a pair was retained in the sample. For the analysis there are 3,708 AD cases, 14,099 controls, and 47,052 AD proxies.

To obtain summary statistics, association analysis was conducted using an approximation of the generalized linear mixed model (GLMM) as implemented in REGENIE (Mbatchou *et al*. 2021) 2.2.4. This multi-stage approach initially fits a whole-genome regression model using ridge regression to estimate polygenic effects while accounting for structure and relatedness (Mbatchou *et al*. 2021). The single variant association model included covariates for sex, age, and five principal components (PCs), which were generated with FlashPCA (Abraham, Qiu, and Inouye 2017) using 153,816 LD pruned variants (r^2^<0.1).

To generate LD reference panels, we computed the LD scores for 378 individuals from 1000 Genomes Project (1kGP) European ancestry using GCTA with the window size was set to 1000 Kb region around a target SNP. LD scores for other populations were computed using 87 Great British (GBR) individuals from 1kGP, following the same methodology used for the EUR (Bulik-Sullivan *et al*. 2015). Additionally, LD scores of genotyped variants were computed for UK Biobank the unrelated subset of 3,708 AD cases and 14,099 controls obtained from UK Biobank. We also obtained the dosage-based LD scores from 420,531 UK Biobank European samples for imputed variants from the Pan-ancestry genetic analysis of the UK Biobank (Pan-UK Biobank) (Karczewski *et al*. 2025).

The h^2^ for AD was estimated using summary statistics and LDSC regression (Tables 1-4) and for comparison using individual-level data and GCTA-GREML (Table 2). For all analyses, age, sex, and five PCs were included as covariates. All h^2^ estimates were computed on a liability scale, assuming a population disease prevalence of 5% (Escott-Price and Hardy 2022). When estimating h^2^ using either GCTA-GREML or LDSC regression, variants with minor allele frequency (MAF) ≤ 0.01, that are strand-ambiguous, or are not SNPs (e.g., insertion-deletions), were excluded. Additionally, only variants that overlap with the LD reference panels were included in the analyses.

**Table 2.** h^2^ estimates for Alzheimer’s disease using UK Biobank data.

| Method | LD reference panel | Sample size - LD reference panel | #Variants | $h^2$ (se) |
| --- | --- | --- | --- | --- |
| GCTA-GREML | - | - | 574,235 | 0.361 (0.039)<sup>a</sup> |
| LDSC regression | UK Biobank study subjects | 17,807 | 574,235 | 0.247 (0.082)<sup>b</sup> |
| LDSC regression | 1kGP GBR | 87 | 502,473 | 0.161 (0.097) |

For all analyses, 3,708 Alzheimer’s disease White British UK Biobank cases and 14,099 White British UK Biobank controls that are unrelated were analyzed. Only genotyped array variants were included in the analysis. $h^2$ was estimated based on a liability of a 5% prevalence in the population. <sup>a</sup>In GCTA-GREML the genetic relationship matrix was built using all 574,235 variants. <sup>b</sup>The LD reference panel was estimated based on the same 3,708 AD cases and 14,099 controls which were used in the analysis. *Abbreviations:* #-number of; 1kGP GBR-1000 Genomes Project Great British; GCTA-GREML-Genome-wide Complex Trait Analysis-Genomic-relatedness-matrix REstricted Maximum Likelihood; LD-linkage disequilibrium; LDSC-linkage disequilibrium score; se-standard error.

## Results

For LDSC regression, h^2^ was highest when estimated using a LD reference panel obtained from the same study subjects which were used to generate the summary statistic. i.e., UK Biobank. This h^2^=0.247 [standard error (se) 0.082] was lower than the h^2^=0.361 (se 0.039) obtained from GCTA-GREML using the same 574,235 variants. For LDSC regression a further underestimation of h^2^=0.161 (se 0.097) occurred when an external LD reference panel (GBR) was used even though it matched the ancestry of the UK Biobank study subjects (Table 3).

**Table 3.** The effect of phenotype definition on h^2^ estimates.

| UK Biobank data analyzed | #Variants | $h^2$ (se) |
| --- | --- | --- |
| 3,708 AD cases & 14,099 controls | 574,235 | 0.247 (0.082) |
| 50,760 cases including 47,052 AD proxies & 14,099 controls | 520,410 | 0.038 (0.024) |
| 47,052 AD proxies & 14,099 controls | 520,391 | -0.006 (0.013)<sup>a</sup> |

For all analyses, summary statistics were generated using UK Biobank study subjects. Only genotyped array variants were included in the analysis. $h^2$ was estimated using UK Biobank in-sample LD reference panel based on a liability of a 5% prevalence in the population. <sup>a</sup>The negative $h^2$ estimate indicates negligible genetic component for this phenotype definition. LDSC regression produces unbounded estimates that can be negative when true $h^2$ is near zero, reflecting sampling variation around a null heritability. *Abbreviations:* #-number of; AD-Alzheimer’s disease; LD-linkage disequilibrium; LDSC-linkage disequilibrium score; se-standard error.

Not surprisingly, the inclusion of proxy AD cases led to a large reduction in the estimated h^2^ from 0.247 (se 0.082) to 0.046 (se 0.023) (Table 3). When the analysis is restricted to controls and proxy AD cases, the h^2^ estimate is further reduced (0.008, se 0.014). We do expect a lower h^2^ for proxy AD cases compared to AD cases since many of the proxies will not develop disease. It should also be noted that the population prevalence for having a first degree relative with AD is ~15% and when this prevalence is used then the h^2^=0.053 (se 0.034), highlighting the impact of using an incorrect prevalence on h^2^ for being a proxy AD case.

For LDSC regression when imputed variants are analyzed, even for those with a high INFO score, there is a decrease in the h^2^ estimate compared to analyzing genotyped variants. When variants have an INFO score between 0.90 and 1 the estimated h^2^ decreases to 0.160 (se 0.041 compared to when genotyped variants are analyzed (h^2^= 0.265, se 0.148) (Table 4). The effect of reducing the estimate of h^2^ when analyzing imputed variants is not limited to LDSC regression. For example, when 744,377 imputed variants with INFO score between 0.90 and 0.92 are included using GCTA-GREML, the estimated h^2^=0.221 (se 0.066), while when genotyped variants were analyzed h^2^=0.361 (se 0.039). Using the same variants with an INFO score between 0.90 and 0.92, LDSC regression provides an estimate of h^2^=0.156 (se 0.350). The median MAF ranges for all groups are around 0.225 (Table 4), suggesting that the underestimation is not due difference in MAFs between imputed and genotyped variants.

**Table 4.** The impact of imputed variants on h^2^ estimates.

| INFO score | Median MAF | #Variants | h<sup>2</sup> (se) |
| --- | --- | --- | --- |
| —<sup>a</sup> | 0.206 | 136,072 | 0.265 (0.148) |
| [0.999,1]<sup>b</sup> | 0.226 | 283,184 | 0.192 (0.072) |
| [0.995,1]<sup>b</sup> | 0.233 | 645,171 | 0.172 (0.050) |
| [0.990,1]<sup>b</sup> | 0.233 | 853,739 | 0.169 (0.044) |
| [0.900,1]<sup>b</sup> | 0.229 | 1,094,814 | 0.160 (0.041) |

For all analyses the summary statistics were generated from 3,708 Alzheimer’s disease White British UK Biobank cases and 14,099 White British UK Biobank controls who are unrelated. h<sup>2</sup> was estimated using UK Biobank LD reference panel, for which only HapMap3 variants (MAF >0.01). h<sup>2</sup> was estimated based on a liability of a 5% prevalence in the population. Only <sup>a</sup>genotyped or <sup>b</sup>imputed variants were analyzed. *Abbreviations:* #-number of; LD-linkage disequilibrium; MAF-minor allele frequency; se-standard error.

## Discussion and conclusion

In this study, we investigated several factors contributing to discrepancies in h^2^ estimates derived from LDSC regression. Our findings reveal significant issues related to mismatched ancestries between summary statistics and LD reference panels, inconsistent phenotype definitions, and the inclusion of imputed genetic variants with varying INFO scores.

Our analysis demonstrates that using LD reference panels with differing ancestries from the summary statistics can lead to substantial inaccuracies in h^2^ estimates. Differences in phenotype definitions, such as including proxies, clinical or autopsied cases, further complicates comparisons across studies. For example, in Jansen et al., 2019 (Jansen *et al*. 2019) (Table 1), AD cases from UK Biobank are defined by clinical diagnosis, self-reported records and family history, whereas Religious Orders Study and Rush Memory and Aging Project (ROSMAP) has autopsy cases. Additionally, the inclusion of imputed variants even those that are imputed with high accuracy, e.g., INFO>0.9 introduces noise, which biases h^2^ estimates downward. These issues emphasize the need for careful consideration of these factors when interpreting h^2^ estimates from LDSC regression.

The AD GWAS meta-analysis in Bellenguez et al., 2022 (Bellenguez *et al*. 2022) included data from 15 European countries, including the Australia, Finland, Greece, Italy, Portugal, and the United Kingdom. Each study within this consortium employs its own case and control definitions, with varying sex and age distributions. Additionally, AD proxies are often included in the meta-analysis in addition to AD cases and controls. The exclusion of AD proxies can lead to a more realistic estimation of AD heritability, e.g., h^2^=0.106 (se 0.012)^24^. When AD proxy cases and AD cases are combined the h^2^ estimates are meaningless because heritability is being estimated for two distinct phenotypes. The low h^2^ estimates in Table 1 can be attributed to several factors, including differences in ancestry between summary statistics, the inclusion of proxy cases, and the use of imputed variants.

A limitation of our analysis is when the number of genotyped variants is low, the standard error of h^2^ estimates becomes relatively high, particularly when the LD reference panel is filtered to only include HapMap3 variants. For example, in Table 4, when only genotyped variants were used to estimate h^2^ and the LD reference panel is limited to HapMap3 variants from UK Biobank, the standard error is 0.148, leading to reduced precision in heritability estimates. This reduces our ability to detect subtle differences in heritability across methods or to make reliable inferences about the genetic architecture of the trait. This often results in overlapping confidence intervals, though the overall trend remains evident. To increase the number of variants, including imputed variants is one approach, but even those with relatively high INFO scores (0.99) can distort h^2^ estimates. Although standard errors of h^2^ can be reduced by increasing the sample size or obtaining summary statistics from meta-analysis of large multiple large studies (Table 1), but this requires careful consideration of potential discrepancies in phenotype definitions and genetic ancestries across cohorts, which can introduce additional biases.

In general, to obtain an unbiased h^2^ estimate, it is ideal to use an in-sample LD reference panel for LDSC regression or to apply methods such as GCTA-GREML, however both of these approaches require individual level data. If individual-level data are not accessible, it is crucial to select an external LD reference panel that matches the ancestry of the summary statistics. Meeting both conditions, matching ancestries for the summary statistics and LD reference panel and using high-quality variant data, ensures that LD patterns are captured with greater precision and that h^2^ estimates are accurate. Therefore, we recommend using only genotyped variants or leveraging sequence data to increase variant count while maintaining high quality. A recent study demonstrated that using whole-genome sequence data and GCTA-GREML largely solved the missing heritability problem for a variety of traits, e.g., standing height, type 2 diabetes and platelet count (Wainschtein *et al*. 2025). It is reasonable to expect that using summary statistics from sequence data and LDSC regression should also lead in an increase in h^2^ estimates.

## Acknowledgements

This study was supported by National Institute of Health [grant number DC017712 (to S.M.L. and A.T.D.)]. We gratefully acknowledge Dr. Longda Jiang from the New York Genome Center and Dr. Jian Yang from Westlake University for their invaluable assistance with the application of GCTA-GREML to analyze imputed variants.

## Web Resources

LD score regression: https://github.com/bulik/ldsc

GCTA: https://yanglab.westlake.edu.cn/software/gcta/

LD scores from UK Biobank: https://pan.ukbb.broadinstitute.org/docs/ld/index.html

1000 Genomes Project data:

https://mathgen.stats.ox.ac.uk/impute/data_download_1000G_phase1_integrated.html

## Data Availability

All summary statistics listed in Table 1 are publicly available. This research also utilized the UK Biobank Resource (application number 32285), a longitudinal cohort of approximately 500,000 individuals aged 40 to 69 years at recruitment between 2006 and 2010. The UK Biobank study received generic approval from the National Health Service’s National Research Ethics Service. The present analyses were approved by the Institutional Review Boards at Yale University (2000026836) and Columbia University (AAAS3494).

## Key points

- LDSC regression heritability estimates can be substantially biased downward when summary statistics are derived from meta-analyses that combine studies with different ancestries, phenotype definitions, or inclusion of proxy cases.
- Including imputed variants in heritability estimation, even those with high imputation accuracy (INFO > 0.9), leads to underestimation of heritability compared to using only genotyped array variants — a bias observed with both LDSC regression and GCTA-GREML.
- Mismatches between the ancestry of the study population and the LD reference panel reduce heritability estimates; using an in-sample LD reference panel yields the highest and most accurate estimates with LDSC regression.
- Combining AD cases with proxy cases in the same analysis produces heritability estimates that are not only lower but effectively uninterpretable, as heritability is being estimated for two distinct phenotypes simultaneously.
- To obtain accurate heritability estimates, researchers should use genotyped or whole-genome sequence variants, ensure ancestry-matched LD reference panels, and apply consistent phenotype definitions across contributing studies.

